# Correlated decision making across multiple phases of olfactory guided search in Drosophila

**DOI:** 10.1101/2020.08.18.256289

**Authors:** Floris van Breugel

**Affiliations:** Dept. of Mechanical Engineering, Graduate Program for Neuroscience, Ecology, Evolution, and Conservation Biology, University of Nevada, Reno, NV 89557

## Abstract

All motile organisms must search for food, often requiring the exploration of heterogeneous environments across a wide range of spatial scales. Recent field and laboratory experiments with the fruit fly, *Drosophila*, have revealed that they employ different strategies across these regimes, including kilometer scale straight-path flights between resource clusters, zig-zagging trajectories to follow odor plumes, and local search on foot after landing. However, little is known about the extent to which experiences in one regime might influence decisions in another. To determine how a flies’ odor plume tracking during flight is related to their behavior after landing, I tracked the behavior of individually labelled fruit flies as they explored an array of three odor emitting, but food-barren, objects. The distance flies travelled on the objects in search of food was correlated with the time elapsed between their visits, suggesting that their in-flight plume tracking and on-foot local search behaviors are interconnected through a lossy memory-like process.

## Introduction

All moving organisms spend a significant amount of their time and energy searching, be it for food, mates, or oviposition and nesting sites. Improving our knowledge of the algorithms that animals use during these search efforts represents a critical step towards understanding how organisms function by connecting neuroscience, behavior, ecology, and evolution [1]. On the behavior and ecology fronts, countless field studies have helped shape our understanding of the search behavior exhibited by mammals, birds, and fish in the context of optimal foraging theory and satisficing [2,3]. In laboratory environments designed to discover the neural basis underlying these decisions, many efforts have focused on olfactory search of organisms including mice [4], insects [5], and crustaceans [6] (for a review, see [7]). To move the field forward, there is a growing push to connect laboratory and field experiments. Perhaps surprisingly, the unassuming fruit fly, *Drosophila melanogaster*, has emerged as a prime model for bridging this gap. Evidence is mounting that despite their numerically simple nervous systems, these creatures are capable of forming visual [8,9] and olfactory memories [10–12], and possess an internal representation of their compass heading with respect to visual cues [13–16]. How these adaptations are involved in search, however, remains an area of active research.

Most natural environments consist of a patchwork of potential resources with a fractal-like distribution, demanding multiple scales of search: long-range, intermediate, local, and nutrient driven. Long-range search for a *Drosophila* consists of flying up to 10 km across the desert to find a new oasis [17,18], initially relying on celestial cues [19,20], as well as vision and wind [21], until it catches an odor plume to follow [22,23]. Within the oasis a fly begins its intermediate search phase: tracking odor plumes [5,24–27] and approaching visual cues [22,28], often relying on the integration of the two to find a fermenting fruit [29,30]. After landing [31], the fly enters its third phase, local search. Now travelling on foot, the fly continues using odors to navigate the patchiness [32–34], as cracks in the skin serve as entry points, whereas mold renders portions too toxic [35]. After tasting some nutrients, the fly enters its final—nutrient driven—search phase, characterized by a so-called “dance” that is largely driven by idiothetic cues [36–38]. Whether or not the fly finds the nutrients it needs, eventually it will decide to take flight and leave, only to start the process all over again. While each of these phases of search has been subject to recent research efforts aimed at understanding both the behavior and neurobiology, little is known about how these individual phases are connected to one another, and how memories from one phase might influence the next.

In this paper I simulate a patchy environment by placing three ethanol-emitting objects in a wind tunnel. Individually marked fruit flies are allowed to freely explore these objects over the course of an 18 hour period. My results indicate that their search behavior on each individual patch is correlated with the time elapsed between patch visits, suggesting that the intermediate and local phases of search are interconnected through a lossy memory-like process.

## Methods

To discover the relationship between decisions flies make during local and intermediate search behaviors in patchy landscapes, I placed three food-barren odorous platforms in a wind tunnel (Fig. 1A-B, [5]). Each platform was set up to constantly emit the attractive odorant ethanol from the center of the platform by bubbling 60 mL/min of air through a 50% ethanol/water solution. The acrylic platforms had a perforated area in the center for emitting the odor, and were all identical in size, shape, color, odor type and concentration. The edge was coated in fluon to prevent the flies from crawling out of view.

**Fig. 1.**
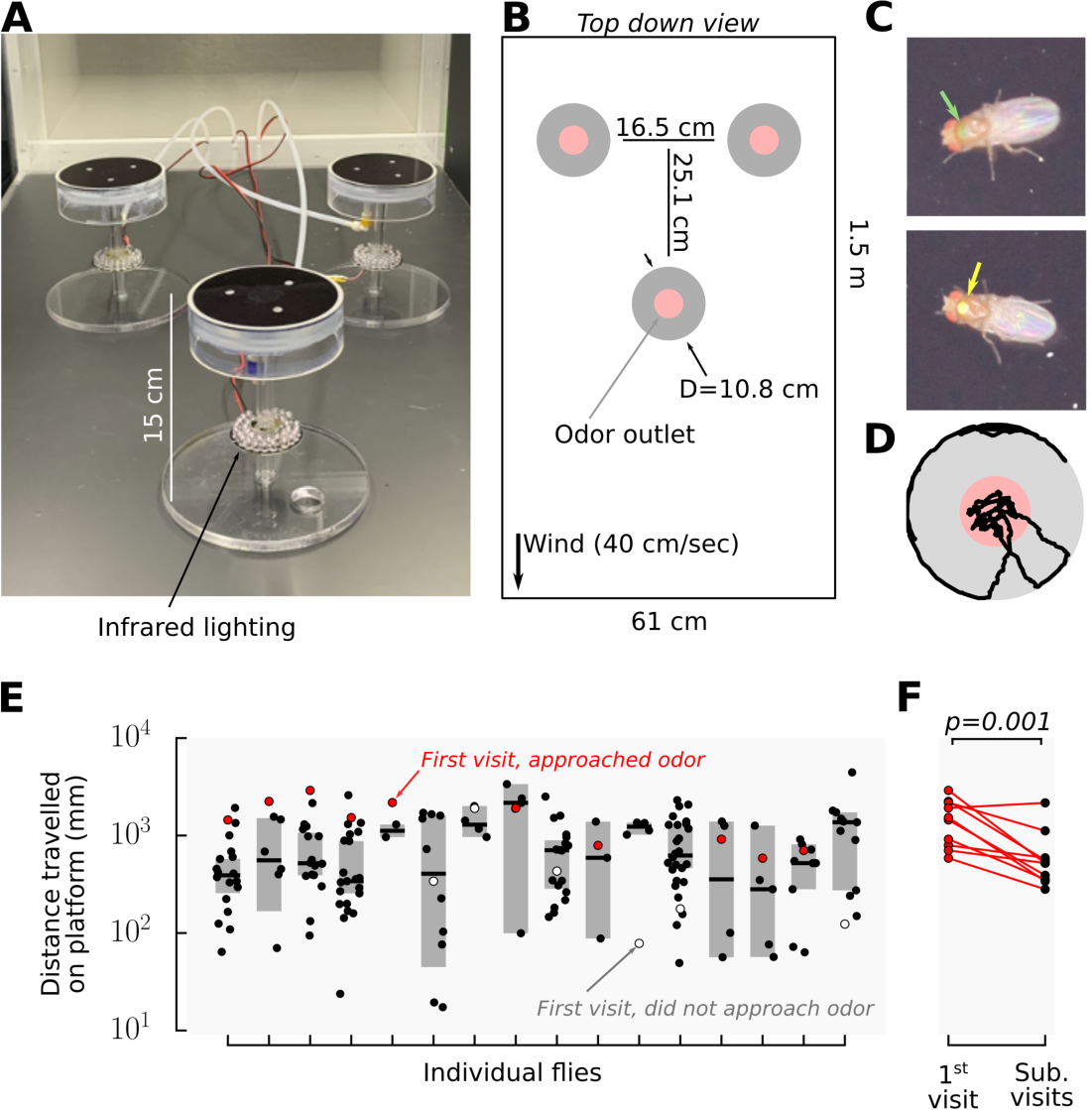
Flies explore odorous objects longer on their first encounter. (A) Photograph of experimental arrangement of three odor emitting platforms inside a 60×60×120 cm^3^ wind tunnel. Above each platform was a machine vision camera to track the flies’ walking trajectories, and a digital SLR to image their markings. (B) Top down diagram of experimental arrangement, red shading indicates approximate region where ethanol odor was emitted, see [5]. (C) Representative photographs of flies indicating their identifying color spots painted on with nail polish. (D) Representative trajectory on one platform. (E) Distance travelled on the platform by individual flies, shading indicates 95% confidence interval about the mean of all visits after the first one. (F) When flies approached the odor on their first visit, they covered more ground than on average during their subsequent visits (paired T-test; t=4.56; p=0.001).

To keep track of individual flies across all three platforms, I painted a dot of colored nail polish on their thorax (Fig. 1C). The flies were cold-anesthetized for the painting, and allowed to recover while being deprived of food, but not water, for 8 hours prior to the experiment start. For each experiment, I used six flies. They were placed in the wind tunnel 6 hours prior to their entrained dusk (relative to a 16:8 light:dark cycle), and allowed to move freely throughout the wind tunnel for 18 hours. When the flies landed on a platform, they were tracked by a machine-vision tracking system described previously [5], with one modification. Every 10 seconds, 18 megapixel color dSLR cameras positioned above each patch photographed the flies. All trajectories were hand-corrected for tracking errors to guarantee their completeness, and manually associated with the correct color identity from the dSLR images.

## Results

In a stereotypical search-bout, flies would spend a significant amount of time near the odor source in the center of the platform, while also making periodic forays towards the edge, often circling the perimeter of the object (Fig. 1D). To quantify their behavior, I focused on the distance the flies travelled during their search, rather than the time they spent as was been done previously [5] (see Discussion for rationale).

The first time the flies’ encountered an odor emitting platform, they walked a significantly larger distance compared to the average of their subsequent encounters, provided that they engaged with the odor during that first encounter (Fig. 1E-F). These results suggest that flies might maintain some form of memory about their failure to find food in order to minimize the time wasted on future fruitless search endeavors. But does this “memory” fade with time? To further investigate this hypothesis, I analyzed their behavior at a more granular scale by comparing the distance travelled to the time elapsed before (pre-interval), and after (post-interval), visiting additional platforms (Fig. 2A).

**Fig. 2.**
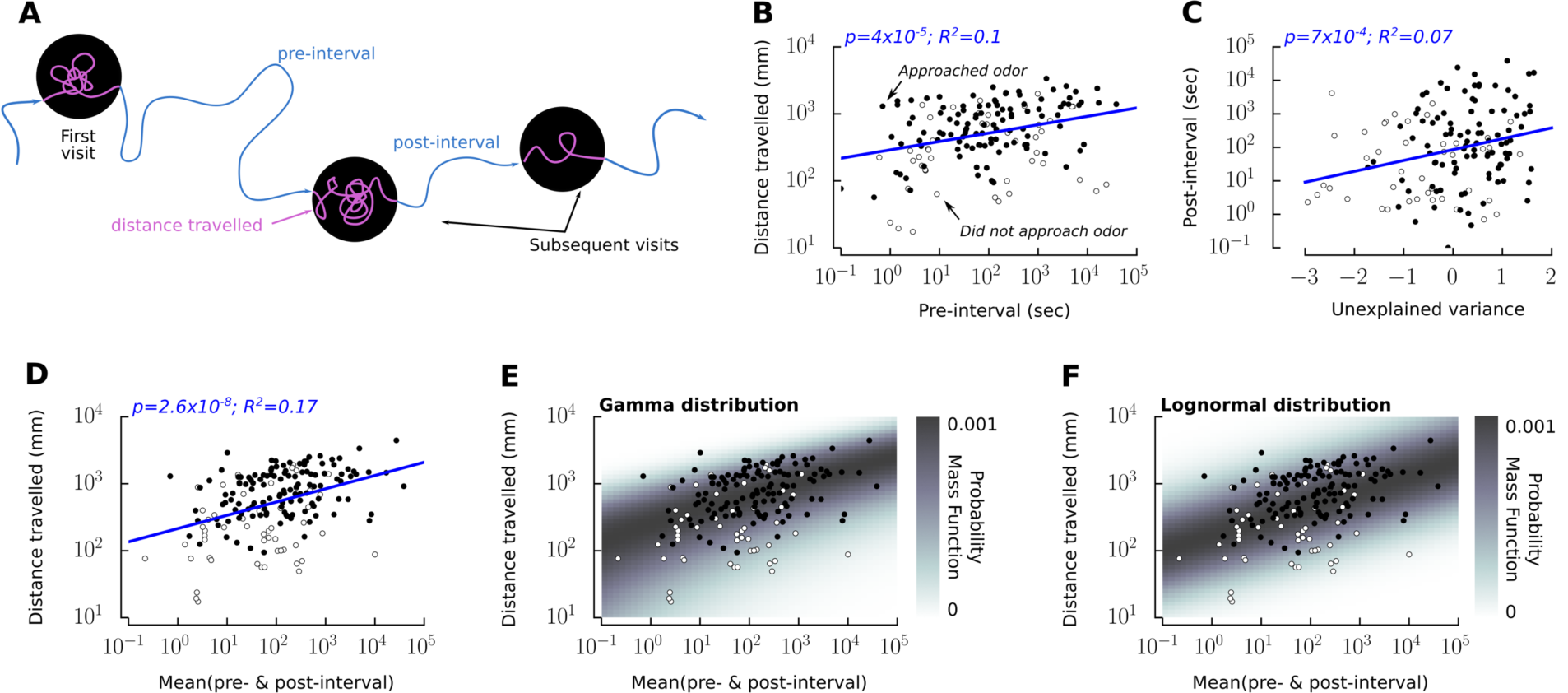
When flies explore an odorous, but food-barren, object, the distance they travel on the object is proportional to the time elapsed since their last visit to a similar object. (A) Cartoon of a hypothetical sequence of visits to three odor emitting patches. (B) Distance flies travel on the platform as a function of the time elapsed since their last visit to the same, or different, platform. Black and white colored dots indicate flies that either approached, or did not approach, the odor during that particular visit, respectively. Blue line shows the linear regression, including all data. Excluding flies that did not approach the odor does not change the statistics in any meaningful way. (C) Post-interval time as a function of the difference between the data and linear regression from B. (D) Relating distance travelled to the mean of the pre- and post-intervals improves the strength of the correlation, plotted as in B. The data in D is well modelled by either a gamma (E), or lognormal (F) distribution, with shape and scale parameters that vary with the abscissa.

Most of the flies explored the odorous patches numerous times, often returning to the same patch several times in a row. Intervals between visits ranged from less than a second, to 16 hours. The distance the flies travelled on the platform while searching is, on a log-log scale, correlated with the time elapsed since they last visited a platform (Fig. 2B). Though significant (p=4×10^−5^), the positive correlation only explains about 10% of the variance (R^2^=0.1). The unexplained variance, however, is positively correlated with the amount of time until their next patch encounter (post-interval) (Fig. 2C). That is, if flies walk for a smaller distance than predicted based on their pre-interval, then their post-interval is likely small as well, allowing them to make up for the missed opportunities. Relating the distance travelled on the platform to the mean of the pre- and post-intervals resulted in a stronger positive correlation than either alone (p=2.6×10^−8^; R^2^=0.17) (Fig. 2D). To ensure that no individual flies played an outsize role in this conclusion, I recalculated the correlation from Fig. 2D in the case where as many as five (32%) random flies were left out of the analysis. The largest p-value for this resampling analysis was 4×10^−5^, indicating that our conclusion is not biased by a small subpopulation.

There was no correlation between distance travelled and time of day (p=0.38, Supp. Fig. 1A), indicating that neither hunger nor circadian rhythm were important factors. Instead, these factors likely determine whether the flies initiate take off and plume tracking. I also did not find a correlation between pre- and post-interval times (p=0.23, Supp. Fig. B), suggesting there are no consistent behavioral sequences such as rapidly flitting between platforms.

## Discussion

My results indicate that search behavior across different search phases—intermediate scale plume tracking and local search on foot—is inter-connected. These observations raise several discussion points: (1) what internal mechanism is responsible for giving rise to the time-interval correlation (e.g. an interval timer, sensory adaptation, or habituation); (2) what mechanism drives flies’ decision to leave a patch; and (3) in what ecological contexts is their behavior advantageous.

### Mechanisms that could give rise to a measure of time-interval

Although a number of models for neural encoding of interval-timing have been proposed [39], there is little experimental evidence for minute to hour scale interval-timing. In rats, extended time sense is encoded in the hippocampus [40–42]. However, the accuracy is modulated by drugs, hormones, and context [43,44]. For insects, the ability to measure time-intervals on the scale of seconds to minutes is open for debate. Parasitoid wasps are capable of learning time intervals [45,46]. Honeybees, however, are not [47,48]. Although their cousins, bumblebees, have been shown to learn fixed time intervals [49], the analysis has been called into question [47]. Instead, their behavior suggests that they may have learned a different strategy that approximates interval timing.

The challenges of reliably encoding time in the nervous system suggest that alternate mechanisms are either wholly, or in part, responsible for giving rise to the observed correlation. One possibility is sensory adaptation, however, this is unlikely given that peripheral olfactory receptor level adaptation occurs on much faster time scales (∼0.5 second [50]) compared to the observed behavior. Instead, habituation is a more likely explanation. Prior experiments with ethanol induced startle responses indicate that habituated responses are attenuated on the time scales of 15-30 minutes [51].

### Mechanisms driving the decision to leave

In ecology, the process of search has been dominated by the field of optimal foraging theory, namely, the Marginal Value Theorem [2] [52] [53] and satisficing [3]. Neither model, however, provides an explanation for the underlying neural mechanisms. A number of “rules of thumb” have been proposed [54,55]. One example is the Threshold Giving Up Time (TGUT) strategy [56] [57]), which states that an animal should continue searching for food in a patch for an amount of time proportional to the quality of the patch. However, because maintaining an accurate sense of time elapsed is challenging, a more likely strategy for insects is to leave after some Threshold Giving Up Distance-travelled (TGUD) has been reached, given the behavioral and neural evidence for insects’ ability to count steps [37,58]. Assuming some variability, a TGUD algorithm should result in a unimodal distribution of distances travelled, but the shape of that distribution could be either normal, or something with a heavier tail such as a lognormal, or levy distribution. An even simpler heuristic for deciding when to leave a patch was proposed in the context of jumping spiders, termed Fixed Probability of Leaving (FPL) [59], where the animal does not need to keep track of time (or distance) and instead leaves with a fixed probability at each time step (or physical step). An FPL strategy would result in an exponential distribution, where the likelihood of leaving after a short distance travelled is very high, falling off exponentially.

Can flies’ decision to leave be modeled by either the TGUD or FPL strategy? A close look at the distribution of distance travelled suggests that although their behavior is not consistent with an exponential distribution it is consistent with both gamma and lognormal distributions, with parameters that vary as a function of the intervals between visits (Fig. 2E-F). At first glance, these results appear to favor the TGUD model, however, a gamma distribution is equivalent to a sum of exponential distributions. Thus, a simple alternative is that the fly has multiple FPL-like processes operating together, which I term Gamma Probability of Leaving (GPL). To distinguish between the TGUD and GPL models will require connecting the behavior with neural activity.

### Ecological context for time-interval correlated search

What advantages might a correlation between intermediate and local search behaviors confer? I propose two hypotheses. First, the fly might form a memory, associating the odor with a low likelihood of finding food, thereby improving search efficiency. However, these relationships might change with the environment and season, so it would make sense for such a memory to revert to an innate value over time. The second hypothesis is that their behavior optimizes search in non-homogeneous environments, where potential resources are clustered in groups separated by larger distances. In this case, it would make sense to tailor search time relative to the ease with which a new potential food source can be found. Distinguishing between the memory and unpredictable environment hypotheses will require experiments that manipulate the memories formed by providing food rewards or manipulating the release of neuromodulators such as dopamine.

**Supp. Fig. 1.**
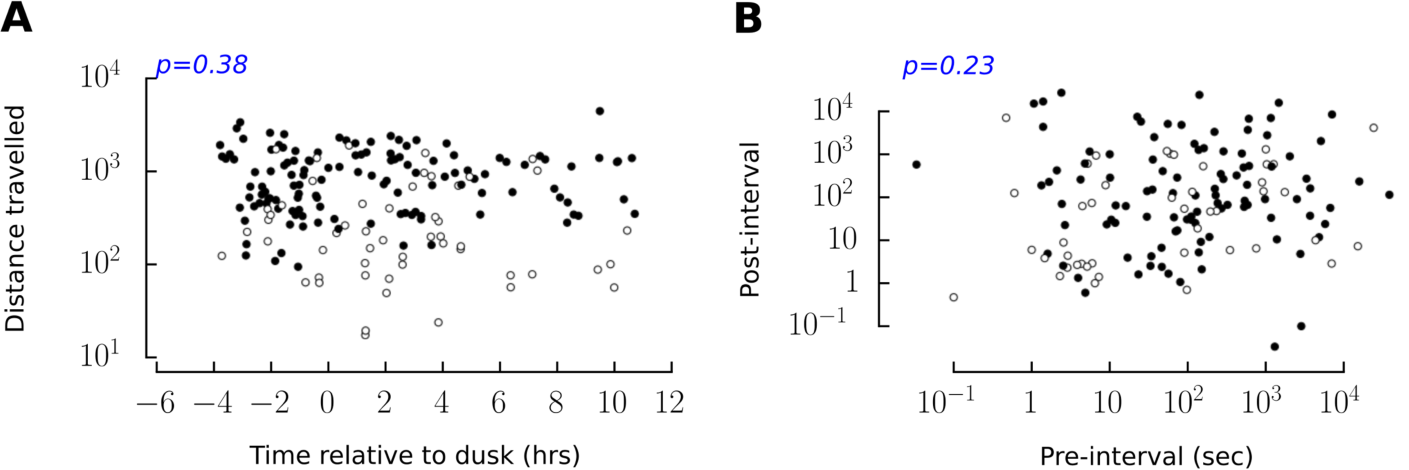
(A) Distance travelled on the platform is not correlated with time of day (p=0.38). (B) Pre- and post-intervals are not correlated with one another (p=0.23). Black and white colored dots indicate flies that either approached, or did not approach the odor during that particular visit.

## Acknowledgements

Many of the ideas presented here originated while I was working in the lab of M H Dickinson, and the experiments were performed in a wind tunnel supported by Jeff Riffell’s National Science Foundation grant (NSF-1626424). I am grateful for the help in analyzing trajectories done by two undergraduates at UNR, Zachary Dugan and Darreann Carmela Hailey, and feedback on the manuscript from Jeff Riffell. The research was supported by a Moore/Sloan Data Science and Washington Research Foundation Innovation in Data Science Postdoctoral Fellowship, a Sackler Scholarship in Biophysics, and the National Institute of General Medical Sciences of the National Institutes of Health (P20GM103650).

